# Correlates of auditory decision making in prefrontal, auditory, and basal lateral amygdala cortical areas

**DOI:** 10.1101/2020.05.14.096263

**Authors:** Julia L. Napoli, Corrie R. Camalier, Anna Leigh Brown, Jessica Jacobs, Mortimer M. Mishkin, Bruno B. Averbeck

## Abstract

Auditory selective listening and decision making underlies important processes, including attending to a single speaker in a crowded room, often referred to as the cocktail party problem. To examine the neural mechanisms underlying these behaviors, we developed a novel auditory selective listening paradigm for monkeys. In this task, monkeys had to detect a difficult to discriminate target embedded in noise when presented in a pre-cued location (either left or right) and ignore it if it was in the opposite location. While the animals carried out the task we recorded neural activity in primary auditory cortex (AC), dorsal lateral prefrontal cortex (dlPFC) and the basal lateral amygdala (BLA), given that these areas have been implicated in auditory decision making, selective listing, and/or reward-guided decision making. There were two main findings in the neural data. First, primary AC encoded the side of the cue and target, and the monkey’s choice, before either dlPFC or the amygdala. The BLA encoded cue and target variables negligibly, but was engaged at the time of the monkey’s choice. Second, decoding analyses suggested that errors followed primarily from a failure to encode the target stimulus in both AC and PFC, but earlier in AC. Thus, AC neural activity is poised to represent the sensory volley and decision making during selective listening before dlPFC, and they both precede activity in BLA.

## Introduction

Spatial selective listening is critical for solving everyday problems including the “cocktail party problem”, which requires attending to one sound source amidst a noisy background of competing sources (Cherry 1953). Common auditory spatial selective listening paradigms in research with humans include modified Posner paradigms in which subjects have to detect auditory stimuli after being cued to a spatial location (Spence and Driver 1994, Alho, Medvedev et al. 1999, McDonald and Ward 1999, Mayer, Harrington et al. 2007, Mayer, Mannell et al. 2009, Roberts, Summerfield et al. 2009, Teshiba, Ling et al. 2013) and selective listening studies (Ahveninen, Huang et al. 2013, Frey, Mainy et al. 2014, Bidet-Caulet, Buchanan et al. 2015).

Previous work in humans has shown that auditory cortex (AC) plays an important role in spatial selective listening tasks, through interactions with prefrontal (Alho, Medvedev et al. 1999) and parietal (Deng, Reinhart et al. 2019) cortex, as well as auditory perceptual decision making (Tsunada, Liu et al. 2016, Christison-Lagay and Cohen 2018). In humans with dorsal lateral prefrontal cortex (dlPFC) damage, studies of selective listening found decreases in a facilitatory evoked activity component, recorded over dlFPC (Bidet-Caulet, Buchanan et al. 2015), and other work has shown that dlPFC supports important aspects of auditory working memory (Plakke, Hwang et al. 2015). It has also been suggested that the basal lateral amygdala (BLA) contributes to attention and value based decision making, in the visual domain, by encoding information about spatial location and the motivational significance of stimuli (Peck and Salzman 2014, Costa, Mitz et al. 2019).

In this work, we describe a spatial selective listening paradigm for monkeys, grounded in tasks used in human studies. In the task, animals first fixate and then press a bar. After this an auditory cue is presented to the left or right side of the monkey. Following the cue there is a delay period, during which white noise is continuously played for added difficulty. At the end of the delay period, a difficult to detect (titrated to about 70% accuracy) target auditory cue is played on the cued or non-cued side, and the animal has to indicate whether the target was on the cued side by releasing the bar, or on the opposite side by not releasing the bar. To investigate the neural correlates of auditory spatial listening, we recorded single unit activity from monkeys as they performed the task. Neural activity was recorded from primary AC, dlPFC, and the BLA. In AC and dlPFC a substantial fraction of neurons were selective to the location of the cue and the subsequent target. We found that AC preceded both dlPFC and BLA in sensory discrimination and also in the decision. Classification analyses of firing rate patterns in error trials indicated that errors during the task were usually the result of a failure to encode the second target stimulus in AC and also in dlPFC. A comparison with a control “passive listening” condition showed that target related activity in dlPFC is almost completely abolished in the passive task, suggesting task-dependent gating of information to areas beyond sensory cortex.

## Methods

The experiments were carried out on two adult male rhesus macaques (Macaca mulatta). The monkeys had access to food 24 hours a day and earned their liquid through task performance on testing days. Monkeys were socially pair housed. All procedures were reviewed and approved by the NIMH Animal Care and Use Committee.

### Experimental Setup

The monkeys were operantly trained to perform a spatial selective listening paradigm. The task was controlled by custom software (Tucker Davis Technologies (TDT) System 3: OpenWorkbench and OpenDeveloper, TDT) which controlled multi-speaker sound delivery and acquired bar presses and eye movements. Eye movements were tracked using an Arrington Viewpoint eye tracking system (Arrington research) sampled at 1 kHz. Monkeys were seated in a primate chair facing a 19-Inch LCD monitor 40 cm from the monkey’s eyes, on which the visual fixation spot was presented. Monkeys performed the task in a darkened, double-walled acoustically isolated sound booth (Industrial Acoustics Company, Bronx, NY). All auditory stimuli were presented from a speaker 10 cm from the left or right of the monkey’s head. Juice rewards were delivered using a solenoid juice delivery system (Crist Instruments).

### Task Design and Stimuli

The monkeys carried out a spatial selective listening task (Fig. 1), modeled after those used in humans. The task required oculomotor fixation throughout the duration of the trial. Both spatial cues and target stimuli were auditory and the monkeys were required to respond when they detected a target embedded in masking noise presented on the cued side. Listening conditions (listen left/right) were blocked with two types of trials (detect/foil) in each condition. At the start of each trial, the monkey was prompted to press a lever and fixate a central point on the screen. After a short delay (2.1-2.4 s), a 50 ms 4 kHz square wave (70 dB) cue was played from a speaker on the left or right of the midline. Frozen diotic white noise (40 dB) was then played from both the left and right speakers from 500 ms after the initial cue until the lever was released. Following a variable delay after noise onset (500, 800 or 1300 ms) a 300 ms 1 KHz square wave target sound was played from either the left or right speaker. If the target sound was on the same side as the cue, it was a detect trial and the animal had to release the lever within 700 ms to receive a juice reward. If the target sound was on the opposite side as the cue, it was a foil trial and the monkey had to continue to hold the lever. Following a second interval of 800 or 1000 ms in foil trials, a second 1 KHz sound was always played on the same side as the original cue. If the animal correctly released the lever following the second target in foil trials it was given a juice reward. Thus, both detect and foil trials were identical in terms of reward expectation. If the response was incorrect, there was a long “timeout” period before the next trial could be initiated. As in our previous work (Camalier, Scarim et al. 2019) the use of square waves (which contain odd harmonics) allowed for wideband stimulation that was perceptually distinct, but whose broad spectral signature robustly activated large swaths of AC in a way that pure tones would not. Thus, similar to human paradigms, the stimuli could be kept identical across all sessions, independent of where recordings were carried out in AC, and data could be collapsed.

**Figure 1.**
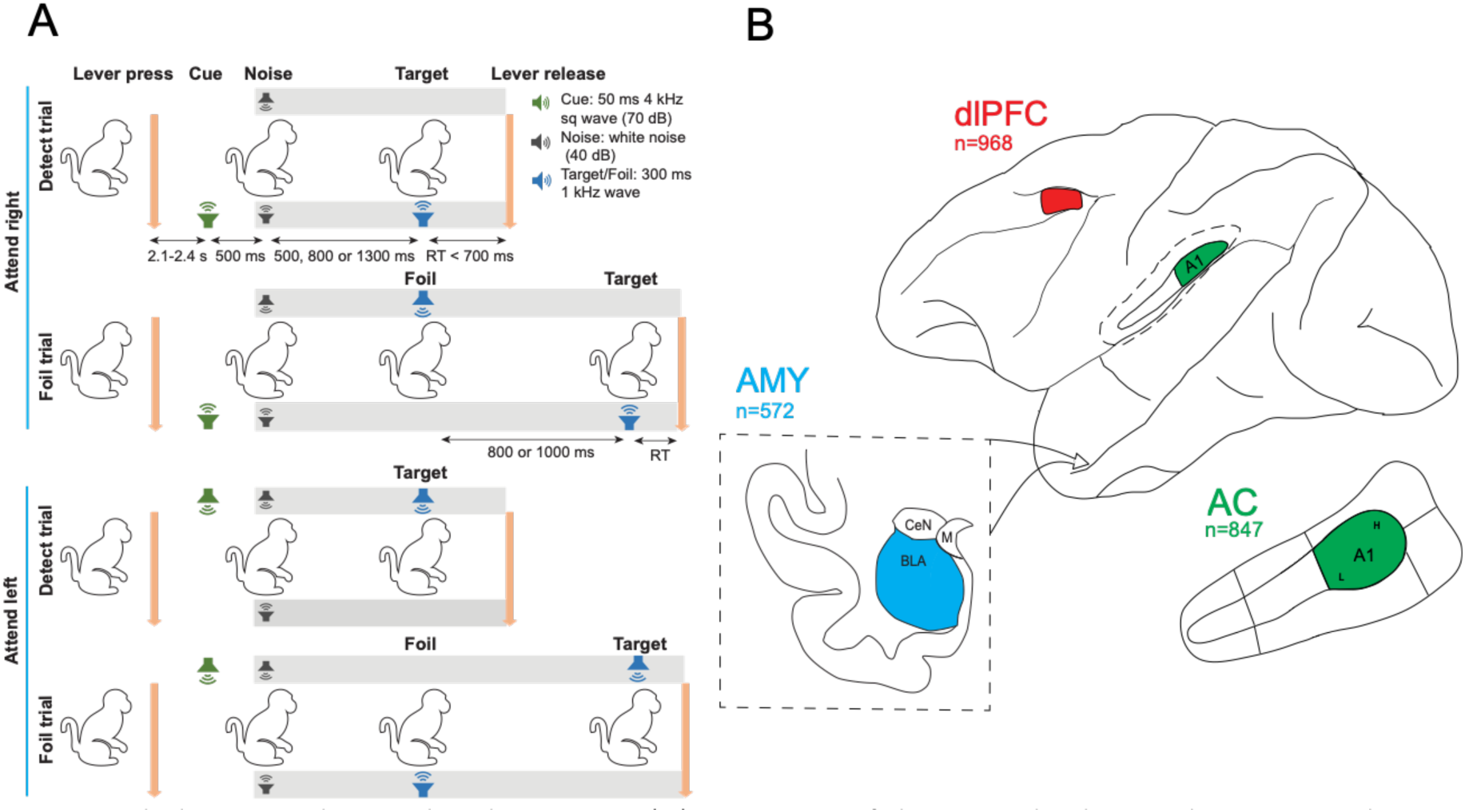
Task design and recording locations. (A) Structure of the spatial selective listening task. Cue conditions (listen left/right) were blocked with two types of trials (detect/foil) in each condition. To begin each trial, the animal must depress a lever and maintain fixation at a central point on the screen. After a short delay (2.1-2.4 s), the animal heard a 4 kHz square wave cue from a speaker on the left or right of its head. A continuous white noise was played 500ms after the initial cue to make target detection difficult. In detect trials, the animal will hear a 1 kHz target (various levels, see Methods) after some stimulus onset asynchrony (SOA) (500, 800 or 1300ms) on the same side as the cue. If the animal releases the lever within 700ms of the target, a fixed juice reward is delivered. In foil trials, the animal will hear a 1 kHz “foil” tone on the opposite side as the cue after the same SOA. The animal must continue to hold down the lever until a 1 kHz target is presented (after 800 or 1000ms) on the same side as the cue. If the animal releases the bar within 700ms of the target, a fixed juice reward is delivered.

The “passive listening” control condition was identical to the active task except the monkey listened passively and did not press a lever, fixate, respond, or receive juice. (B) Recording locations of single neurons across auditory cortex (AC), dorsolateral prefrontal cortex (dlPFC) and the basal lateral amygdala (BLA). (Top) Patch of dlPFC recording area morphed to anatomical landmarks. (Middle) AC grid coverage on region A1 of auditory cortex based on topography, latency and frequency reversals. (Bottom) Region of interest highlighted in blue--the entire left basolateral amygdala--targeted by v-trodes. In all three areas, we selectively recorded from the left hemisphere of the animal.

Several psychometric quality controls were included to ensure that the monkeys were consistently performing the task across sessions. The sound level of the target for the two monkeys was individually titrated to maintain performance at ∼70% correct. Thus, the detection was difficult. For monkey 1, target tones were delivered at levels between 20-30 dB, with the majority of tones in the 20-22 dB range. For monkey 2 target tones were delivered at levels between 30-40 dB, with the majority of tones in the 33-36 dB range. Within a session, the target sound varied 0-3 dB from trial to trial to ensure that the monkeys were responding to the target side and not to consistencies in (or guessing based on) sound level differences between speakers that may have resulted from otherwise undetectable differences in calibration between the two speakers. Periodic “catch trials” (∼10%) with a 0-dB target tone were included to ensure that the monkeys were responding to the target and not timing relative to the presentation of the cue or noise. In addition, we included easy trials (∼10%) with the target sound played at 40dB. These trials helped maintain motivation and ensured that the monkey could perform the experiment at 100% accuracy when sufficient sensory information was available. Similarly, to encourage motivation during foil trials (which lasted longer), the “detect/bar release” target after a foil sound was louder than typical target sounds for each monkey (monkey 1: 27 or 30 dB; monkey 2: 35 or 40dB).

Before the task was run, a battery of passive listening and mapping stimuli were played. Within this battery was a control condition of “passive listening”. In this condition the monkey was presented with the task stimuli with trial types and timing matched to the selective listening task. However, the animals did not press or release a bar, fixate, or receive juice rewards. This task allowed us to compare sensory responses between active and passive task conditions. Monkeys were cued that this was a passive condition as they did not have access to the lever or juice tube, and it was done as part of a passive-listening mapping battery, consistently before the start of the active task.

### Neurophysiological Recordings

Monkeys were implanted with titanium headposts for head restraint before data collection began. Custom acrylic chambers were designed and fitted to the monkeys in a separate procedure to allow vertical grid access to the left dorsolateral prefrontal cortex (Fig. 1B; dorsal bank of the principal sulcus extending ventral, >1mm away from arcuate sulcus, roughly 46/8Ad), the basal and lateral portions of the amygdala (entire dorsoventral extent), and auditory cortex (primarily A1 but including small portions of lateral belt areas). Recording areas were verified through a T1 scan of grid coverage with respect to underlying anatomical landmarks, combined with maps of frequency reversals and response latencies of single neurons to determine A1 location and extent (Camalier, D’Angelo et al. 2012, Camalier, Scarim et al. 2019). Recordings were mainly carried out in simultaneous AC and PFC sessions, with BLA sessions occurring later in the experiment, but some data included were from just one, or even three simultaneously recorded areas in a given session. We recorded the activity of 2,387 single neurons during the task (N = 847 (AC), N = 968 (dlPFC), and N = 572 (BLA) across monkeys 1 and 2).

In both monkeys, we recorded using either 16 or 24 channel laminar “V-trodes” (Plexon, Inc, Dallas TX; 200-300μm contact spacing, respectively). The electrodes allowed for identification of white matter tracts, further allowing identification of electrode location with respect to sulci and gyri. Electrodes were advanced to their target location (NAN microdrives, Nazareth, Israel) and allowed to settle for at least 1 hour before recording. Neural activity was recorded either primarily simultaneously (AC and PFC) or primarily individually (BLA), although there were some sessions in which all 3 areas were recorded from.

Multichannel spike and local field potential recordings were acquired with a 64-channel Tucker Davis Technology data acquisition system. Spike signals were amplified, filtered (0.3-8kHz), and digitized at ∼24.4 kHz. Spikes were initially sorted online on all-channels using real-time window discrimination.

Digitized spike waveforms and timestamps of stimulus events were saved for sorting offline (Plexon sorter V 3.3.5). Units were graded according to isolation quality (single or multiunit neurons). Single and multiunit recordings were analyzed separately, but patterns were similar, so they were combined. The acquisition software interfaced directly with the stimulus delivery system and both systems were controlled by custom software (OpenWorkbench and OpenDeveloper, controlling a RZ2, RX8, Tucker Davis Technologies (TDT) System 3, Alachua, FL).

### Data analysis

For the ANOVA and PSTH analysis, all trials on which monkeys released the lever in the correct interval were analyzed. Trials in which the monkey answered incorrectly (∼30% of all trials), were excluded. The average number of correct trials analyzed for the ANOVA and PSTH analyses were 467.05 (AC: 480.11, dlPFC: 471.72, and BLA: 439.81 trials). We performed a 2 x 2 ANOVA (cue x target) on the activity of single neurons. The response is given by the interaction in this ANOVA. The dependent variable was the firing rates of individual neurons. All trials in which the monkeys correctly released the lever within 700ms of the target were analyzed. The firing rate of each cell was computed in 300 ms bins advanced in 25 ms increments. We separated the analysis into three different segments of time, locked to the time surrounding the individual presentations of the cue, noise, and first target.

Next, we created a population post-stimulus time histogram (PSTH) for the firing rates of the individual neurons with respect to cue condition (left/right) and trial condition (detect/foil). For this analysis the firing rate of each cell was computed in 1 ms bins and smoothed with a 3 bin moving average. Data are plotted using 25 ms bins, but t-tests, to determine onset latencies, were computed on the 1 ms bins.

For the decoding analyses, we separately analyzed correct and error trials. A trial was considered correct if the monkey released the lever after the presentation of the appropriate target within 700 ms. All other trials were deemed incorrect. The average number of error trials analyzed for decoding was 123.53 (AC: 127.57, dlPFC: 136.99, and BLA: 94.77). For neural analysis, the firing rate of each cell was computed in 100 ms bins and advanced in 25 ms increments. Decoding analyses were performed using leave-one-out cross-validation to predict which observations belong to each trial condition using the SVM classifier in Matlab. All decoding was done using pseudo-populations composed of all neurons recorded from a structure across all sessions. Trials were assigned randomly from the different sessions within each condition.

For the decoding analyses, we calculated significant differences between correct and error trials using a bootstrap analysis (Efron and Tibshirani 1998). We generated data according the null hypothesis that there were no differences between correct and error trials. We did this by sampling with replacement, from the combined set of correct and error trials, sets of bootstrap correct and error trials. Both the null correct and error bootstrap sets contained combinations of correct and error trials. We then carried out the decoding analysis using the bootstrap trials to determine the decoding accuracy when correct and error trials were mixed. We did this 1000 times. We calculated the difference in fraction correct between correct and error trials in each time bin, for each set of bootstrap trials. This gave us 1000 differences sampled from the null distribution, between correct and error trials, in each time bin. We then compared the difference in the actual data to the differences in the null distribution, and computed a p-value, which was the relative rank of the true difference in the null distribution samples. That is to say, if the true difference was larger than, for example, 986 samples in the null distribution, it was significant with a two sided p-value of 2x(1000-986)/1000 = 0.028.

## Results

### Task, recordings, and single cell encoding of task factors

We recorded neural activity from 2 monkeys while they carried out a spatial selective listening task (Fig. 1A). At the start of each trial, the monkeys acquired central fixation (Fig. 1A), and pressed a bar. After a baseline hold period, an auditory stimulus (the cue) was presented from a speaker on the left or right of the monkey. After the cue there was a delay period during white noise was played, continuing until bar release. Following the delay period, a second stimulus (the target) was presented on the same or opposite side as the cue. The monkeys were trained to release the bar if the cue and target were on the same side (detect trials) and continue to hold if they were not on the same side (foil trials). In foil trials, following a second delay after the first target sound, a second target sound was played that was always on the same side. In detect trials the mean response time was 374.9 ms (std = 27 ms).

While the animals carried out the task, neural activity was recorded (Fig. 1B), from three areas: auditory cortex (AC, N = 847), dorsal lateral prefrontal cortex (dlPFC, N = 968) and the basal lateral amygdala (BLA, N = 572). We began by carrying out ANOVA analyses on correct trials, for each single neuron. With the ANOVA we examined the effects of cue location, target location, and their interaction (which determines the response), using spike counts in a 300 ms window, advanced by 25 ms (Fig. 2). During the cue period, we found that activity discriminated cues rapidly in AC (Fig. 2A). In dlPFC, activity discriminated cues as well, but the effect increased slowly (Fig 2D). The BLA, however, showed minimal cue discriminative activity, with the number of neurons coding cue location only slightly above chance (Fig. 2G). Note that cue location was blocked in the task, which led to small baseline, statistically significant, elevation of cue side encoding prior to cue presentation. Although encoding peaked in AC and dlPFC following the cue, elevated cue discrimination was maintained during the delay interval, which was not affected by the white noise, in both AC and dlPFC. The BLA showed less delay period activity.

**Figure 2.**
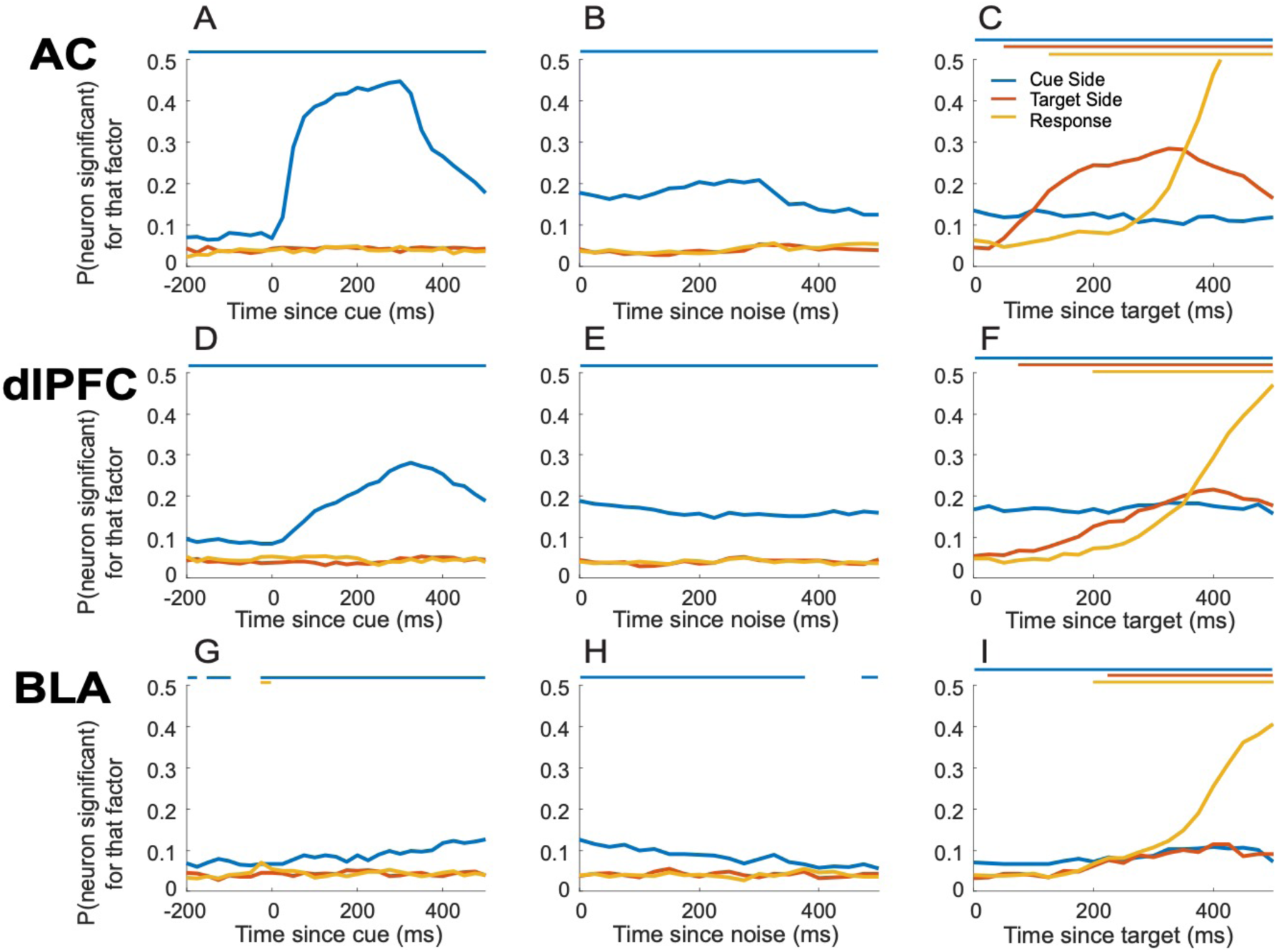
ANOVA analysis. Recording of single neurons from caudal AC (A1, lateral belt), dlPFC and BLA while monkeys are performing a spatial selective listening task. A 2 x 2 factor ANOVA (Cue side x Target side, p < 0.05) using 300ms bins sliding at 25ms. Bin endpoint was used to align time on the x-axis, i.e. 300 ms is a bin from 0 to 300 ms. Bars above each plot represent the bins in which a statistically significant fraction of neurons encode each factor by color. (A, D, G) During presentation of the cue, neurons respond differentially to the cue location. (B, E, H) Post-cue, a substantial fraction of neurons is selective to cue side, during the delay period, in both AC and dlPFC. (C, F, I) Post-target presentation, a substantial portion of neurons in all three areas of interest are selective to the response.

When the target stimulus was presented, it was rapidly and robustly encoded in AC (Fig. 2C). The dlPFC also encoded the target stimulus (Fig. 2F), although later than AC, which would be expected. The BLA only weakly encoded the target stimulus and only at about the time of the response (Fig. 2I). The response was also encoded first in AC (Fig. 2C), after which it was encoded in dlPFC (Fig. 2F). The response, unlike the cue and target locations, was robustly encoded in the BLA (Fig. 2I). Overall, all variables, including the delay period activity and the response, were encoded first and most robustly by AC. The dlPFC did encode all task factors, but after AC. The BLA showed only weak encoding of the cue and the target but robustly encoded the response.

To calculate sensitive statistics on potentially low firing rate neurons the ANOVA analysis used relatively large time windows. These time windows, however, do not allow determination of precise onset times for task factors. To characterize onset times at a finer time scale, we calculated PSTHs using 1 ms time windows, smoothed with a 3-point moving average, for each neuron (Fig. 3 – plotted using 25 ms bins). We then carried out t-tests (p < 0.01, uncorrected) in each bin to estimate the time at which the population in each area discriminated between conditions. We found that the cue was discriminated in AC at 26 ms (Fig. 3B) and in dlPFC at 78 ms (Fig. 3D) after stimulus onset. Using these small bins, the population of BLA neurons did not discriminate cue side, likely due to low firing rates (Fig. 3F). The target was discriminated in AC at 37 ms (Fig. 3G), in dlPFC at 211ms (Fig. 3I) and in the BLA at 274ms (Fig. 3K) after tone onset. Finally, the decision was discriminated in AC at 278 ms (Fig. 3H), in dlPFC at 334 ms (Fig. 3J) and in BLA at 330 ms (Fig. 3L) after target onset. Therefore, AC precedes both dlPFC and BLA in sensory discrimination and decision discrimination at the time of target presentation, and this result was consistent, with two analysis approaches at 2 different time scales.

**Figure 3.**
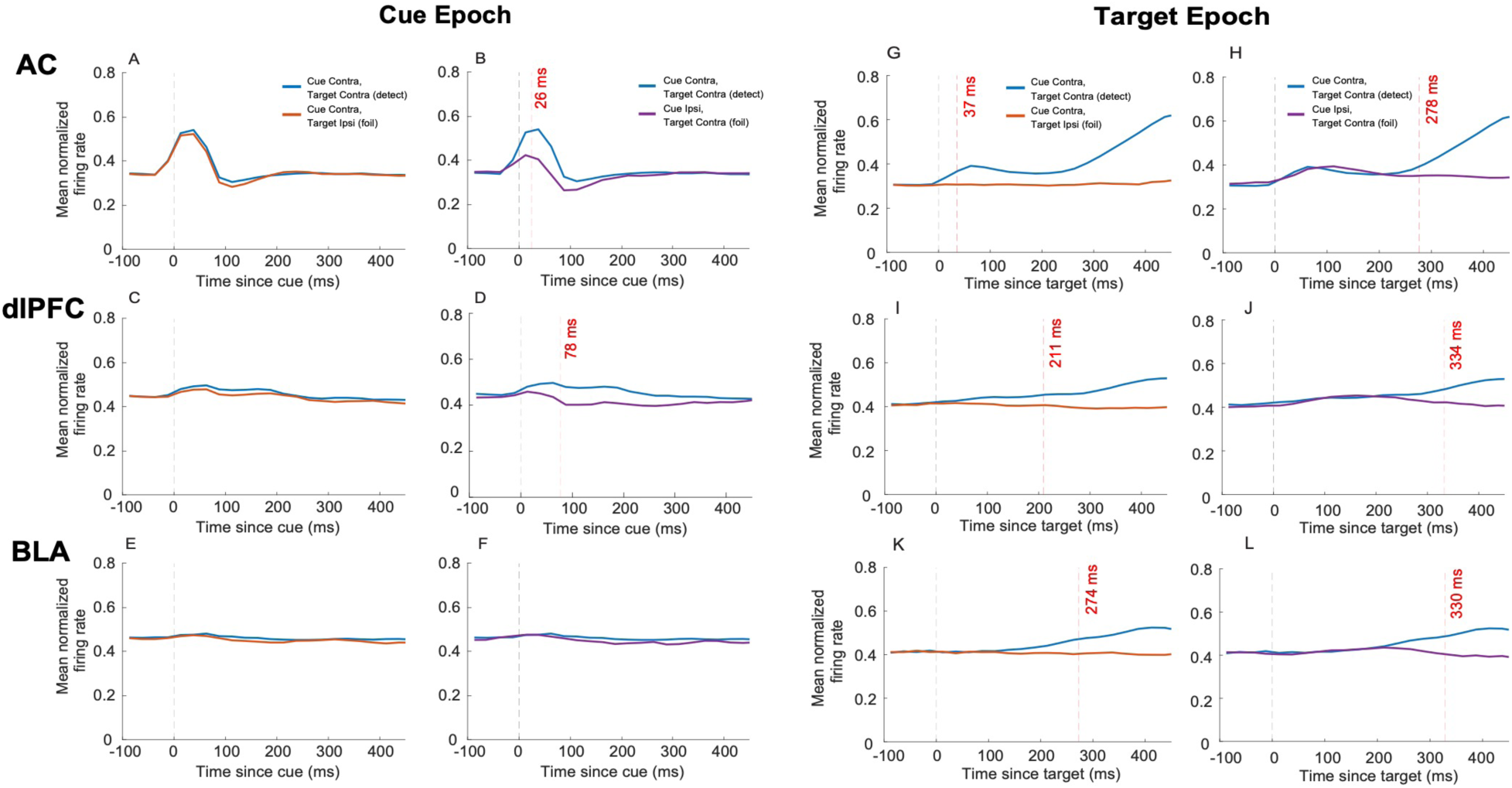
Post-stimulus time histograms (PSTHs). Mean normalized firing rates of neurons plotted using non-overlapping 25ms bins smoothed with a 3-point moving average. Bin midpoint was used to align time on the x-axis. Analysis was conducted to assess precise timing of changes in neuronal firing rates in AC, dlPFC and BLA. Paired t-tests were performed on all bins to determine significant difference in firing rates. (A, C, E) compares conditions that are identical in cue side but vary in target side, as a measure of sensory identification. (B, D, F) compares conditions that are identical in target side but vary in the cue location (left or right). (G, I, K) Conditions are matched for cue side but vary in target side. (H, J, L) Conditions shown have opposite cue sides but matched target side.

### Decoding correct and error trials

In the next analyses we used decoding to examine error trial activity. We were interested in which processes broke down in error trials. To examine this, we used leave-one-out cross validation on pseudo populations (see methods) to predict, using the neural activity, the side on which the cue was presented (Fig. 4), the target was presented (Fig. 5) and the response (Fig. 6). The decoding model was first estimated using only correct trials. We then classified the error trials using the decoding model estimated on correct trials, to see if neural activity in error trials represented the stimuli that were presented, and the response that was made. We found that the neural population in both AC and dlPFC rapidly predicted the cue location (Fig. 4A, D), and maintained prediction through the delay interval (Fig. 4B, D), consistent with the single-neuron results. The BLA did not discriminate clearly the cue side (Fig. 4G). There were no significant differences between correct and error trials for cue encoding, and this finding was consistent through the delay interval. Therefore, the cue was correctly encoded in error trials.

**Figure 4.**
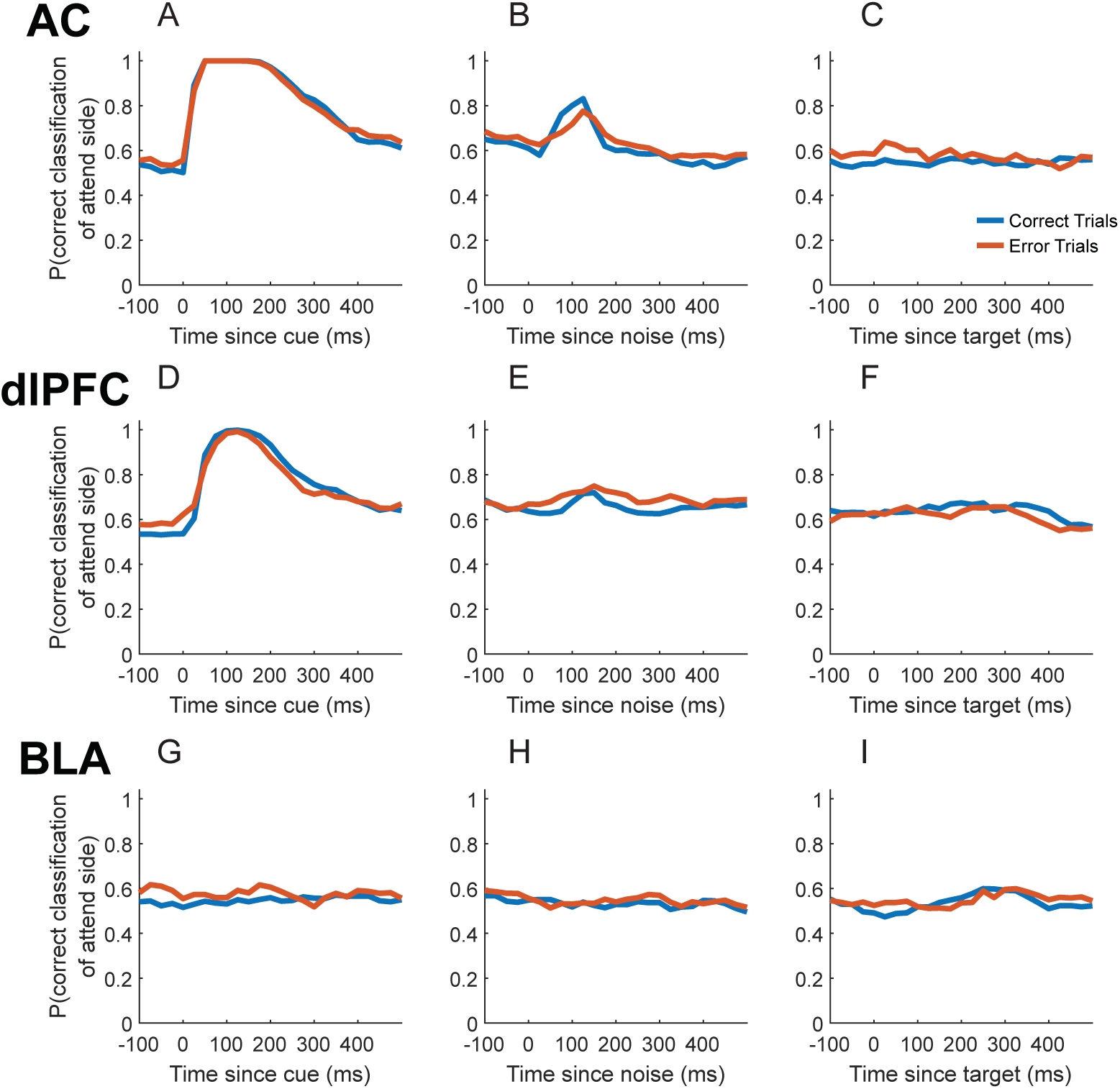
Classification analysis to cue location factor comparing correct and error trials. Analysis was performed using 100 ms bins, sliding at 25 ms. Bin endpoint was used to align time on the x-axis. Analysis performed using leave-one-out cross-validation to predict which observations belong to each cue condition. Bootstrap test performed with 1000 pseudorandom samples. No significant difference in classification rates was found between correct and error trials in any brain region during any time bin.

**Figure 5.**
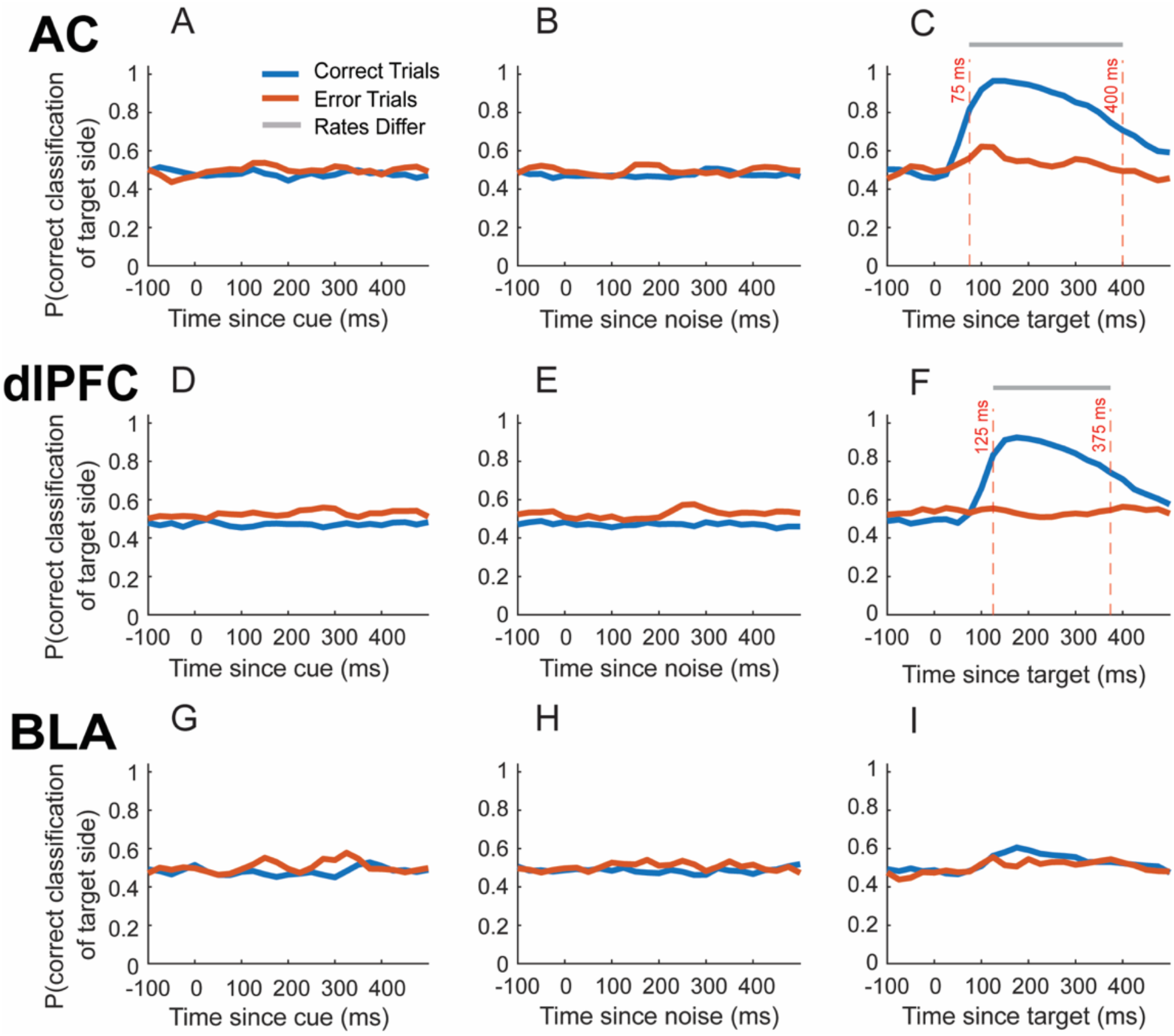
Classification analysis to target factor by correct or error. Analysis was performed using 100 ms bins, sliding at 25 ms. Bin endpoint was used to align time on the x-axis. Grey shaded areas represent timepoints where correct and error classification rates differ (p < 0.01, bootstrap). (C) In AC, a significant difference is seen in classification rates during the target epoch that begins after 75ms and ends after 400ms. (F) In dlPFC, the difference in classification rates begins slightly later and ends slightly earlier, starting at 125ms post-target and ending at 375ms post-target.

**Figure 6.**
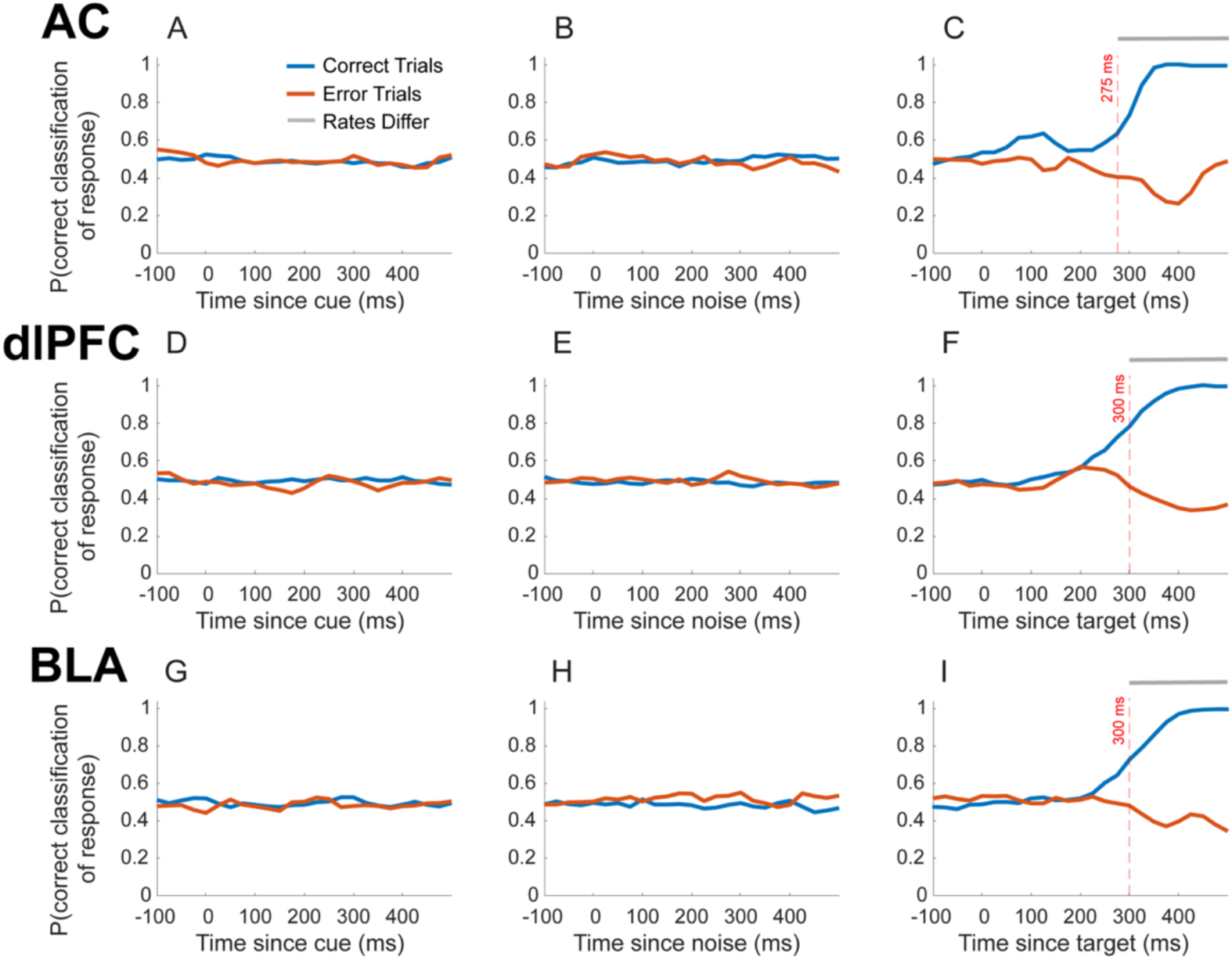
Classification analysis to response. Grey shaded areas represent timepoints where correct and error classification rates differ. Bin endpoint was used to align time on the x-axis. (C) Differences between correct and error trials in AC were from (275ms to 500ms), (F) In dlPFC from (300ms to 500ms) and (I) In BLA from (300ms to 500ms).

When we decoded the target side using neural activity, we found that in correct trials the target location was robustly predicted by AC (Fig. 5C) and dlPFC (Fig. 5F). There was minimal prediction of the target in the BLA (Fig. 5I). In error trials, however, the target was not well predicted by any of the areas (Fig. 5). The correct and error trial predictions diverged (p < 0.01 bootstrap) 75 ms after target onset in AC and 125 ms after target onset in dlPFC.

In error trials, animals either released when they should not have, or did not release when they should have. When we predicted the response, relative to what the monkeys should have done, we found an accurate prediction in correct trials in all 3 areas (Fig. 6C, 6F, 6I). Furthermore, in error trials, the neural activity in all 3 areas predicted the response the animal actually made, and not the response they should have made, because the prediction of what the monkeys should have done went below 0.5. Consistent with the other analyses, we found that predictions in error and correct trials diverged statistically in auditory cortex (275 ms after target onset) and subsequently in dlPFC and BLA (300 ms after target onset).

Next, we examined the position of the population neural activity relative to the discrimination boundary, extracted from the decoding model. For the decoding analysis (Fig. 4-6), this quantity is thresholded in each trial and time-bin, and the time-bin in that trial is classified as either, e.g. cue left or cue right, depending on whether the position is positive or negative. However, the average distance to the decoding boundary provides a continuous estimate of how well the population discriminated the conditions vs. time (Fig. 7). In general, these analyses were consistent with the thresholded decoding analysis. Cue related activity diverged in correct and error trials, reflecting the cued side, and the activity in error trials matched the activity in correct trials (Fig. 7A, D, G). The breakdown in activity following target presentation could also be seen (Fig. 7B, E, H). The response encoding dynamics did reflect the fact that the wrong response tended to be predicted by population activity (Fig. 7C, F, I). However, it could be seen that the activity diverged less than it did in correct trials, consistent with the lower decoding performance.

**Figure 7.**
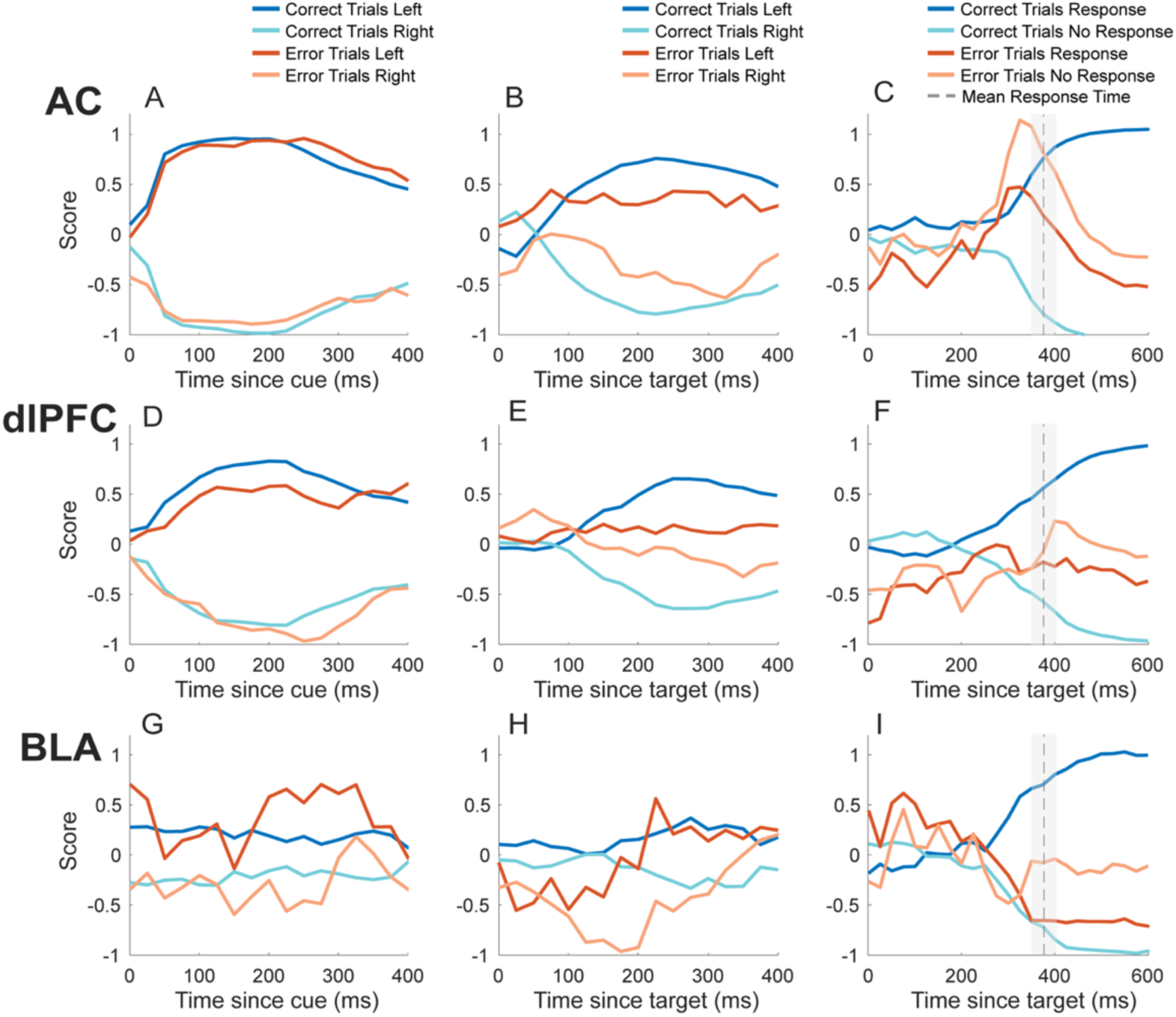
Distance to classification boundary derived from support vector machine classifier. Analyses were conducted using 200ms bins sliding at 25ms. Bin endpoint was used to align to the x-axis. Error trials were defined as trials in which the animal responded incorrectly or responded outside of the allotted reaction time window. Correct trials are presented in shades of blue and error trials in shades of red. The reaction time for detect trials is shown as a dotted line, with the standard deviation shaded, in panels C, F and I. (A, D, G) Conditions were separated by whether a trial was cued on the left or right side of the animal. (B, E, H) Conditions were separated by whether the first target was presented on the left or right side of the animal. (C, F, I) Conditions were separated by whether there was a response or no response made by the animal, i.e. if it was a detect or foil trial, respectively. For error trials, conditions were separated by whether there should have been a response or no response, regardless of what the animal chose to do.

In a final series of analyses, we analyzed data from a passive task, collected in each session before the main, active task data. The sensory stimulation in the passive task was identical to the stimulation in the active task, except the animals did not press a bar to initiate a trial, they did not release the bar to indicate their choice, and there was no juice tube so they could not be rewarded. When we examined encoding of cue location, we again found robust coding in AC (Fig. 8A). All of the other signals, however, were much weaker. The cue responses in dlPFC dropped from a peak near 30% in the active task to about 10% in the passive task (Fig. 8D). Interestingly, there was delay activity in the passive task, in AC (Fig. 8B), perhaps because the animals were highly over-trained. The delay activity in dlPFC was reduced from about 20% of the population to about 10% (Fig. 8E). There was also a small amount of target encoding in AC (Fig. 8C). Target encoding in dlPFC did not exceed chance (Fig. 8F). Encoding in the BLA only sporadically exceeded chance, perhaps due to type-I errors, or low-level encoding (Fig. 8G-I).

**Figure 8.**
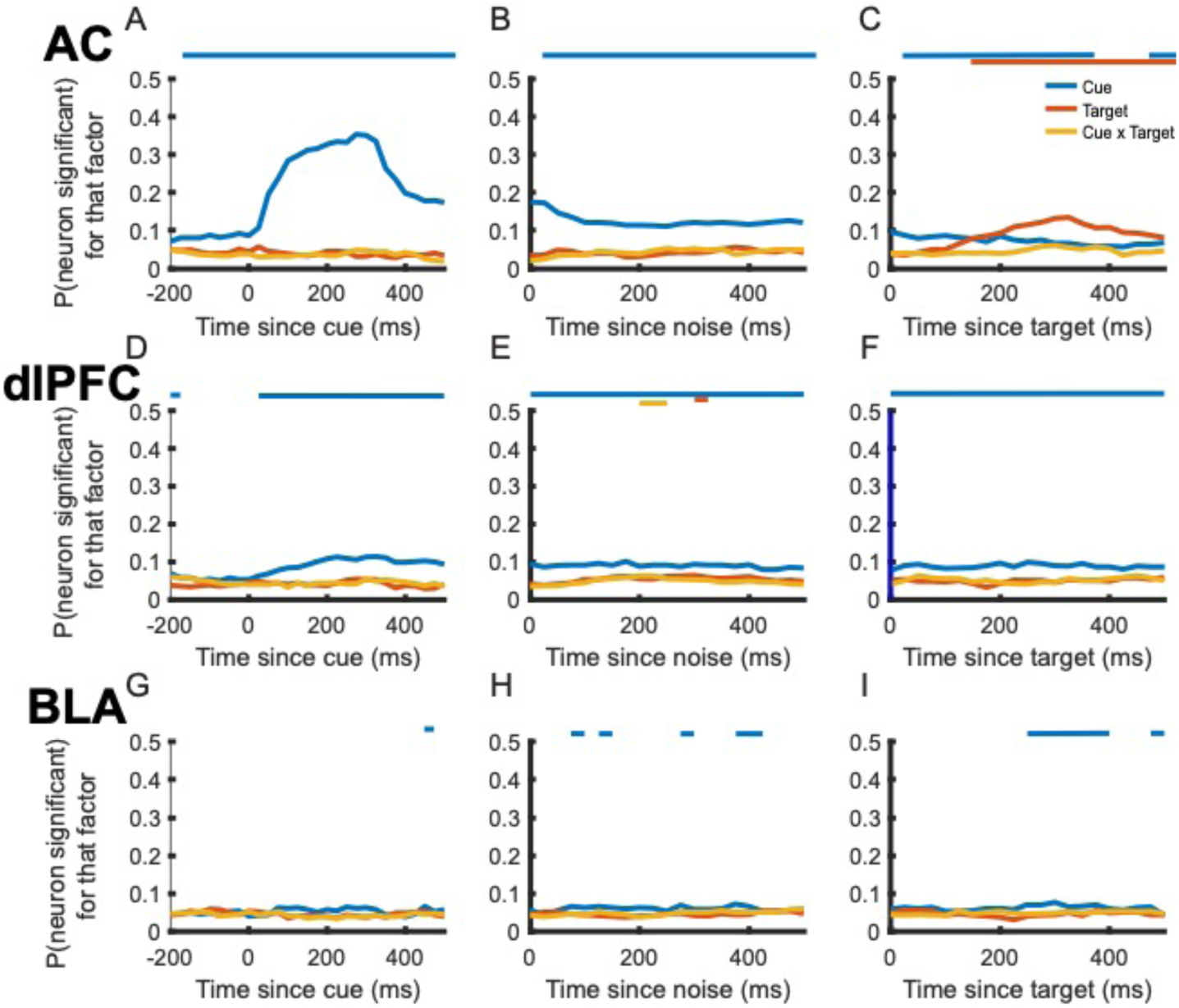
Passive task data. ANOVA analysis of data from passive condition, which was identical to the task, except the monkey was simply required to sit passively and listen to the task structure’s sounds. Bars above each plot represent the bins in which a statistically significant fraction of neurons encode each factor by color. (A, D, G) During presentation of the cue, neurons respond differentially to the cue location. (B, E, H) Post-cue, neurons are selective to cue side, during the delay period, in both AC and dlPFC, but weakly, compared to the active task. (C, F, I) Post-target presentation, only AC encodes the target, and none of the areas encode the response.

We also examined onset times, using small time-bins (Fig. 9). We found differences in responses in AC that depended on the side of the stimulus for the cue at 32 ms (Fig. 9B) and for the target at 53 ms (Fig. 9G). However, we did not detect population level differences in responses, using these small time bins, in dlPFC or BLA, which suggests that responses that reached significance in the ANOVA analyses were driven by low firing rates. Overall, beyond cue encoding in AC, responses across all 3 areas were reduced in the passive task, relative to the active task.

**Figure 9.**
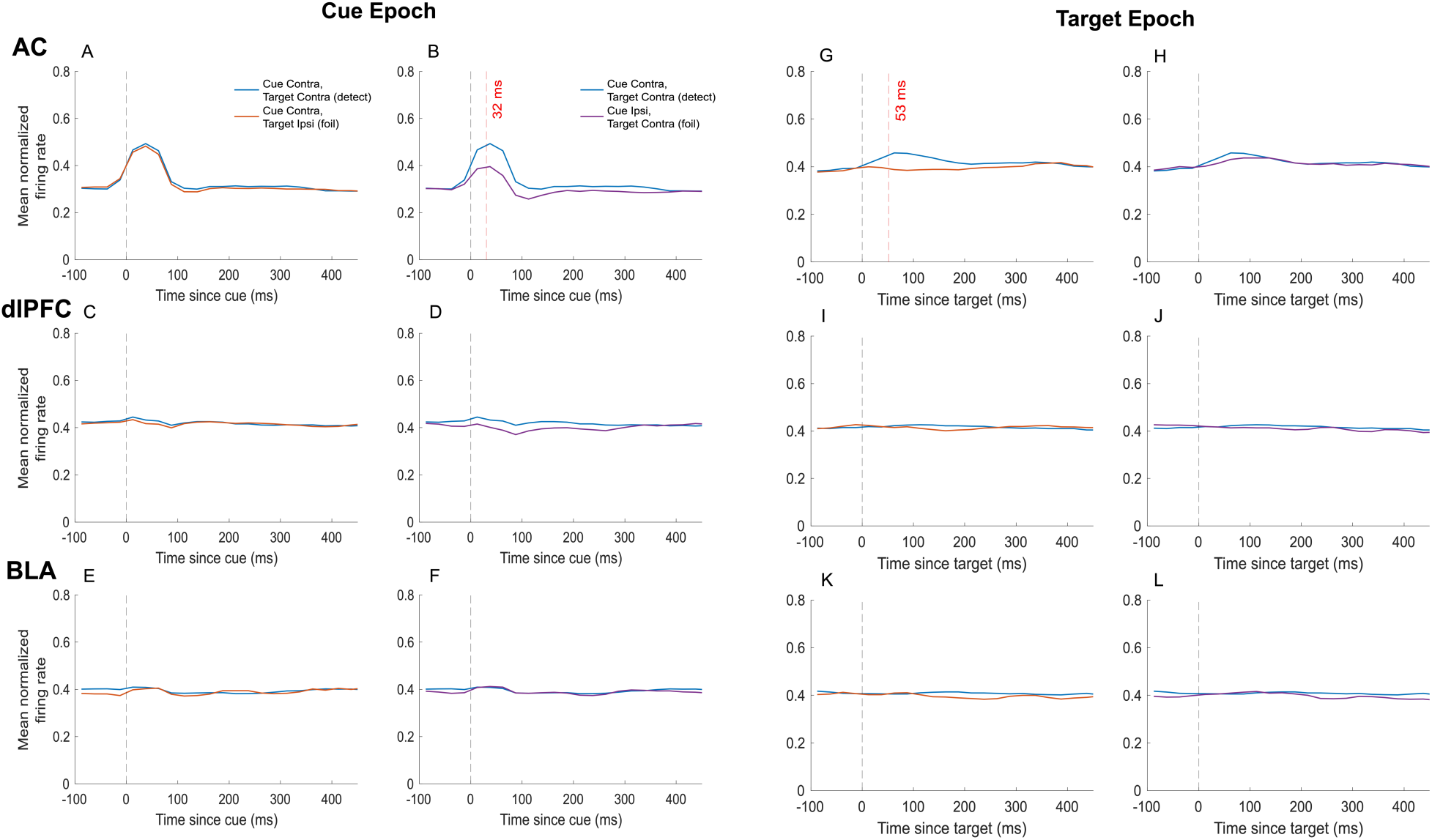
Passive data. Post-stimulus time histograms (PSTHs). Mean normalized firing rates of neurons plotted using non-overlapping 25ms bins smoothed with a 3-point moving average. Bin midpoint was used to align time on the x-axis. Analysis was conducted to assess precise timing of changes in neuronal firing rates in AC, dlPFC and BLA. Paired t-tests were performed on all bins to determine significant difference in firing rates. (A, C, E) compares conditions that are identical in cue side but vary in target side, as a measure of sensory identification. (B, D, F) compares conditions that are identical in target side but vary in the cue location (left or right). (G, I, K) Conditions are matched for cue side but vary in target side. (H, J, L) Conditions shown have opposite cue sides but matched target side.

## Discussion

We trained monkeys on a selective listening task, based on tasks used in humans. The task required animals to detect a difficult to discriminate auditory stimulus, embedded in white noise. We found that AC encoded cues, targets, and decisions, prior to either dlPFC or the BLA. In addition, AC had delay activity that coded the location of the initial cue. It is not clear, however, whether the AC delay activity depended on dlPFC delay activity, or even parietal activity that we did not record. Activity in dlPFC closely followed activity in AC. The BLA, on the other hand, only minimally encoded cue and target activity. The BLA was strongly engaged, however, at the time of choice, although the choice related activity followed activity in AC. Therefore, the AC appears to support many of the functions required for auditory selective listening. This is in contrast to early visual areas, which represent visual features, but play a minimal role in decision making aspects of tasks (Britten, Shadlen et al. 1992, Zaksas and Pasternak 2006).

Previous work has shown that AC neurons can encode non-sensory, choice-related activity (Huang, Heil et al. 2019). The Huang et al. study found that whether a response was predictable following a cue tone, based on the task condition, affected neural responses in AC to the tone. Therefore, AC encoded whether the response was determined by the first cue. Our results are consistent with this and other studies (Christison-Lagay and Cohen 2018), in that we show that auditory cortex encodes the necessary response. However, in our task, the response was not determined by the first cue, so response related activity only followed the target. We also show, that encoding in AC precedes encoding in dlPFC, and we dissociated through our fully crossed experimental design, encoding of cue location, target location, and the required response. Although it is possible that AC inherits response encoding from a cortical area other than dlPFC, the anatomical organization of this system suggests it would have to be a nearby area, for example belt or parabelt auditory cortex (Romanski and Averbeck 2009, Kajikawa, Frey et al. 2015, Tsunada, Liu et al. 2016), or perhaps the medial geniculate. Given that AC is deeper into the neural processing stream than, for example, primary visual cortex (Mizrahi, Shalev et al. 2014), it seems possible that AC could have sufficiently sophisticated mechanisms to compute the required response locally. Though AC precedes PFC in the encoding of the decision in both correct and error trials, the responses across areas are also quite similar within this task (∼50 ms differences). When responses during the passive condition were analyzed, the fraction of responsive neurons was reduced and responses were later in all areas relative to the task-related responses (and BLA was completely unresponsive, consistent with a primary role in reward guided behavior). Particularly, dlPFC showed a reduction of responses to the cue and delay activity and an abolishment of target related activity compared to the active task condition. This is consistent with data from the same animals and areas during a passive oddball task in which dlPFC activity was later (∼100 ms) and weaker than in AC (Camalier, Scarim et al. 2019). Taken together it suggests that the strength and timing of the information transfer between AC and dlPFC can be flexibly allocated and is dependent on task demands. Lastly, comparison of the active and passive conditions highlights the sustained nonsensory motor/reward related activity in “primary” sensory cortex (AC)(Knyazeva, Selezneva et al. 2020).

Both prefrontal (Green, Doesburg et al. 2011, Bidet-Caulet, Buchanan et al. 2015) and parietal (Michalka, Rosen et al. 2016, Deng, Reinhart et al. 2019, Deng, Choi et al. 2020) cortex have been shown to play important roles in auditory spatial attention in humans. Although we found clear responses related to the cued side in dlPFC, they followed AC. This was true of not only the sensory responses, but also the decision response. From our data it is not, however, possible to determine whether the delay period activity, which may represent sustained attention/working memory for the cue location, was sustained by AC, dlPFC, or their interaction. In addition, several of the spatial attention paradigms used in the human work required participants to attend or discriminate sounds in one location, while ignoring sounds on the contralateral side (Deng, Reinhart et al. 2019). It is possible that if we had required the monkeys to carry out complex perceptual discriminations at one location, while ignoring distractors at another location, we would have found stronger engagement of dlPFC. We did use a white masking noise, following the cue signal, to examine its effects on behavior and neural representations of the cue location. Although we did see some effects of the noise onset in the decoding analysis, effects which were stronger in AC than dlPFC, they were transient and resulted in increased decoding accuracy for the cued location. The increased accuracy may have followed from an overall increase in neural activity, which may have improved decoding performance. Also, we did not record neural activity in parietal cortex, which may also play a role in the sustained delay period activity, although it would be interesting to consider inferior parietal cortex in future studies.

We found that the BLA played little role in encoding the cue location, and responses related to the choice followed responses in AC. This is inconsistent with previous reports of the BLA’s involvement in visual-spatial attention (Peck, Lau et al. 2013). In these tasks, the amygdala neurons encoded the valence of stimuli, that were saccade targets, during delay periods (Peck and Salzman 2014). There are several differences between these tasks, and ours, however. For example, the tasks used in Peck et al. were based on visual-spatial paradigms instead of an auditory-spatial paradigm, and they also required eye movements to spatial locations. Although the BLA receives auditory inputs (Romanski and LeDoux 1992, Romanski and LeDoux 1993), these inputs may play a smaller role in the primate than they do in rodents (Munoz-Lopez, Mohedano-Moriano et al. 2010). In rodents, auditory cues can be associated with shock in Pavlovian fear conditioning (Romanski and LeDoux 1992). These studies have shown that the amygdala plays an important role in the associative process between cues and shocks. Although, the amygdala is also involved in reward guided behavior (Costa, Dal Monte et al. 2016, Averbeck and Costa 2017, Costa, Mitz et al. 2019). We did find a small, although significant, population of amygdala neurons, that encoded the auditory cue and the auditory target. They did so, however, at long latencies. Therefore, the BLA appears to play a minimal role in the cognitive process of selective listening under reward-constant trials in highly trained animals.

The present study also shows a substantial dissociation of function between the BLA and dlPFC. This dissociation differs from the similarity between these structures seen in reinforcement learning (RL) tasks, in which both dlPFC and the BLA show substantial encoding of the identity of visual stimuli, the reward values associated with those stimuli, and reward outcomes (Bartolo, Saunders et al. 2019, Costa, Mitz et al. 2019). The primary difference between the BLA and dlPFC, in RL tasks, is that the dlPFC strongly encodes the direction of eye movements required to saccade to a rewarding visual stimulus (Bartolo, Saunders et al. 2019), whereas the BLA encodes eye movement directions only at a low level (Costa, Mitz et al. 2019). Thus, in RL tasks, the BLA and dlPFC show similar responses, which are also similar to those seen in the ventral striatum (Costa, Mitz et al. 2019) and orbito-frontal cortex (Costa and Averbeck 2020), with which the BLA is mono-synaptically connected. The current study, however, shows that in cognitive, auditory selective listening tasks, the BLA and dlPFC show different responses, until the animal makes a reward guided choice.

## Conclusions

We found that AC encoded cues, targets, and decisions, before dlPFC, in an auditory selective listening task. We also found that AC had delay period activity. The BLA had minimal cue or target activity, although it did encode decision activity. The decision related activity in the BLA, however, followed decision related activity in AC. Overall, this suggests that AC may carry out most important computations relevant to auditory selective listening. The main caveat is that it is not possible to determine whether delay period activity, which likely critically underlies performance in this task, is supported by AC in the absence of dlPFC or parietal cortex. Future work, for example inactivating dlPFC and/or parietal cortex (Plakke, Hwang et al. 2015), while recording in AC, could clarify this question.

## Acknowledgements

The authors would like to thank Dr. Richard Krauzlis (NEI/NIH), Dr. Barbara Shinn-Cunningham (Carnegie Mellon) and Dr. Brian Scott (NIMH/NIH) for valuable input on task design and training, Dr. Richard Saunders (NIMH/NIH) for surgical assistance, and the NIH Section on Instrumentation for assisting in custom manufacture of recording chambers and grid. Research was supported by Intramural Research Program of the National Institute of Mental Health NIMH DIRP ZIA MH002928-01 to BA and ZIA MH001101-25 to MM.

